# Evolution of Environmentally-Enforced, Repeat Protein Topology in the Outer Membrane

**DOI:** 10.1101/309393

**Authors:** Meghan Whitney Franklin, Sergey Nepomnyachiy, Ryan Feehan, Nir Ben-Tal, Rachel Kolodny, Joanna S.G. Slusky

## Abstract

Outer membrane beta barrels (OMBBs) are the proteins on the surface of Gram negative bacteria. These proteins have diverse functions but only a single topology, the beta barrel. It has been suggested that this common fold is a repeat protein with the repeating unit of a beta hairpin. By grouping structurally solved OMBBs by sequence, a detailed evolutionary story unfolds. A strand-number based pathway manifests with progression from a primordial 8-stranded barrel to 16-stranded and then to 18-stranded barrels. The transitions from 16- to 18-stranded barrels show mechanisms of strand number variation without domain duplication, such as a loop to hairpin transition. This indicates that repeat protein topology can be perpetuated without genetic duplication likely because the topology is being enforced by the membrane environment. Moreover, we find the evolutionary trace is particularly prominent in the C-terminal half of OMBBs which may be relevant to understanding OMBB folding pathways.

## Introduction

Outer membrane proteins have a remarkably homogeneous topology. All known Gram negative, outer membrane proteins, save one (Dong, et al. 2006), are right-handed, up-down beta barrels with the N and C termini of the barrel remaining on the membrane face from which they inserted. Yet, these proteins carry out all the functions necessary for the interface between the cell and its environment: adhesion, various specific and nonspecific forms of import and efflux, pilus formation, and proteolysis.

The two main structural differences that modulate function among OMBBs are the barrel’s girth and the location of functional amino acids. The barrel’s girth is a function of strand number. Bacterial OMBBs each have an even number of strands between 8 and 26. The addition of each hairpin (two strands connected by a loop) imparts an extra 1.66Å to the radius (Franklin, et al. 2018) in the same way that the addition of an extra two planks to a pickle barrel widens the pickle barrel’s radius by a consistent amount. Change in radius of a barrel alters pore specificity to accommodate or reject molecules for passage based on size. Widening or narrowing the barrel also changes the location of the loops to make for better adhesion or pilus formation.

Function can also be diversified by locations of particular amino acids. Diversity of amino acids allows similarly sized pores to have different chemical specificity. Moreover, chemically potent residues impart proteolytic capabilities.

Although there are few instances in which the membrane barrel topology was converged upon independently (Franklin, et al. 2018), most OMBBs are homologous as shown by sequence alignments (Remmert, et al. 2010; Reddy and Saier 2016). Like all diversification events, the diversification that maintained this common fold while accommodating so many diverse functions occurred as a combination of amplification, recombination, and the accretion of mutations.

The placement of functional amino acids is most commonly attributed to the accretion of mutations. Conversely, that the OMBBs are all composed of different numbers of tandem hairpins (two strands connected by a loop) is the basis of early studies demonstrating that OMBBs are a form of repeat protein (Neuwald, et al. 1995). Recently it has been clarified that the original repeat unit of the beta hairpin or double hairpin has been amplified to create barrels of different strand numbers (Remmert, et al. 2010)

In general, repeat proteins are believed to be generated by duplication and recombination within a single gene and less commonly by polymerase or strand slippage of DNA hairpins (Marcotte, et al. 1999). Such gene and motif duplication is a common mechanism for generating biological diversity and has occurred through evolution resulting in larger proteins and in increasing the diversity of protein function (Ohno 1970; Zhang 2003). Gene duplication is known to be especially prevalent in membrane proteins (Shimizu, et al. 2004). However, the lack of exons in prokaryotes eliminates the possibility of complex editing and recombination events as a mechanism of increasing biological diversity.

The diversification of the OMBBs depends in part on the relative rates of duplication and mutations. Though the relative rates of these two pathways vary by organism, overall duplications are more common than substitution mutations (Katju and Bergthorsson 2013). Mutation rate in *E. coli* is 5.4 × 10^−10^ substitutions per base pair per replication (Drake, et al. 1998), while duplication rate in the closely related *Salmonella* is of 2 × 10^−3^ to 4.6 × 10^−6^ duplications per gene per generation (Reams, et al. 2010). At approximately 1×10^3^ base pairs per OMBB protein, the rate of a protein acquiring a single mutation is indeed less than the rate at which a protein will acquire a duplication. However, which of the mutations or duplications are fixed is determined by concerns of the benefit, detriment, or neutrality of the diversification event.

Here we consider the OMBB path of diversification by tracing sequence alignments among their solved structures. Although many studies exclusively use sequence data to understand the relationships among proteins, here the use of structural data allows us to determine how genetic changes influence structural changes in previously undocumented ways. Using structural data we find a strand number based pathway of evolution from 8-stranded barrels to 16-stranded barrels to 18-stranded barrels. Moreover, we find evidence that strand number diversification can occur through mutation, complicating the longstanding view that repeat topology arises from repeating DNA. Finally, we find that the C-terminal half of OMBBs show particularly strong evolutionary trace signal, which may be relevant to understanding the OMBB folding pathway.

## Results

### Several groups of OMBBs

We used the data set described previously (Franklin, et al. 2018) composed of 130 structurally characterized OMBB proteins which are <85% sequence-similar to each other, including 113 which are less than 50% similar to each other. These include single OMBBs, multi-chain OMBBs, and multi-chain lysins. Hidden Markov Model (HMM) profile alignments of our dataset were generated to determine evolutionary relationship. HMMs determine homology using a probabilistic model with the sequences as the inputs. The results were given as sequence alignments and a score called an E-value which is the expected number of false positives in the database with a score at least as good as that of the match (Soding 2005) These results were mapped into a network model with proteins as nodes and alignments as connections drawn based on the magnitude of the E-value. An interactive version of the network, allowing visual inspection of the underlying sequence and structural similarity, is available online (http://cytostruct.info/rachel/barrels/index.html

It is difficult to determine what E-value cut off to use for assessing OMBB homology. Though the usual rule of thumb is to only use E-values less than 10^−3^ (Pearson 2013), bacterial evolution has been extremely long and fast, giving it approximately 10^12^ opportunities for introducing genetic variation such as amplification, recombination and accretion of mutations. In addition, membrane proteins are less conserved than soluble proteins (Sojo, et al. 2016) and the lipid-facing side even more so (Jimenez-Morales and Liang 2011). Here, our highest E-values cut off is 10^−2^ though we often show the data for lower E-values as well. The interactive, online network has a scroll bar so that the user can change the E-value cut off at will.

In this manuscript we focus exclusively on the largest group which we call the prototypical beta-barrels. The large majority (73% or 95/130) of OMBBs fall into this single highly-interconnected group (see Table S1 for the list of PDBs used). Our group of structurally solved outer membrane beta barrels is composed of multiple subclusters (fig. 1 and supplemental GIF). The subclusters are organized by strand number, with most barrels that share a strand number being more closely related than barrels that differ in the number of strands. The relationships between the subclusters articulate evolutionary relationships between the strand numbers.

**Figure 1.**
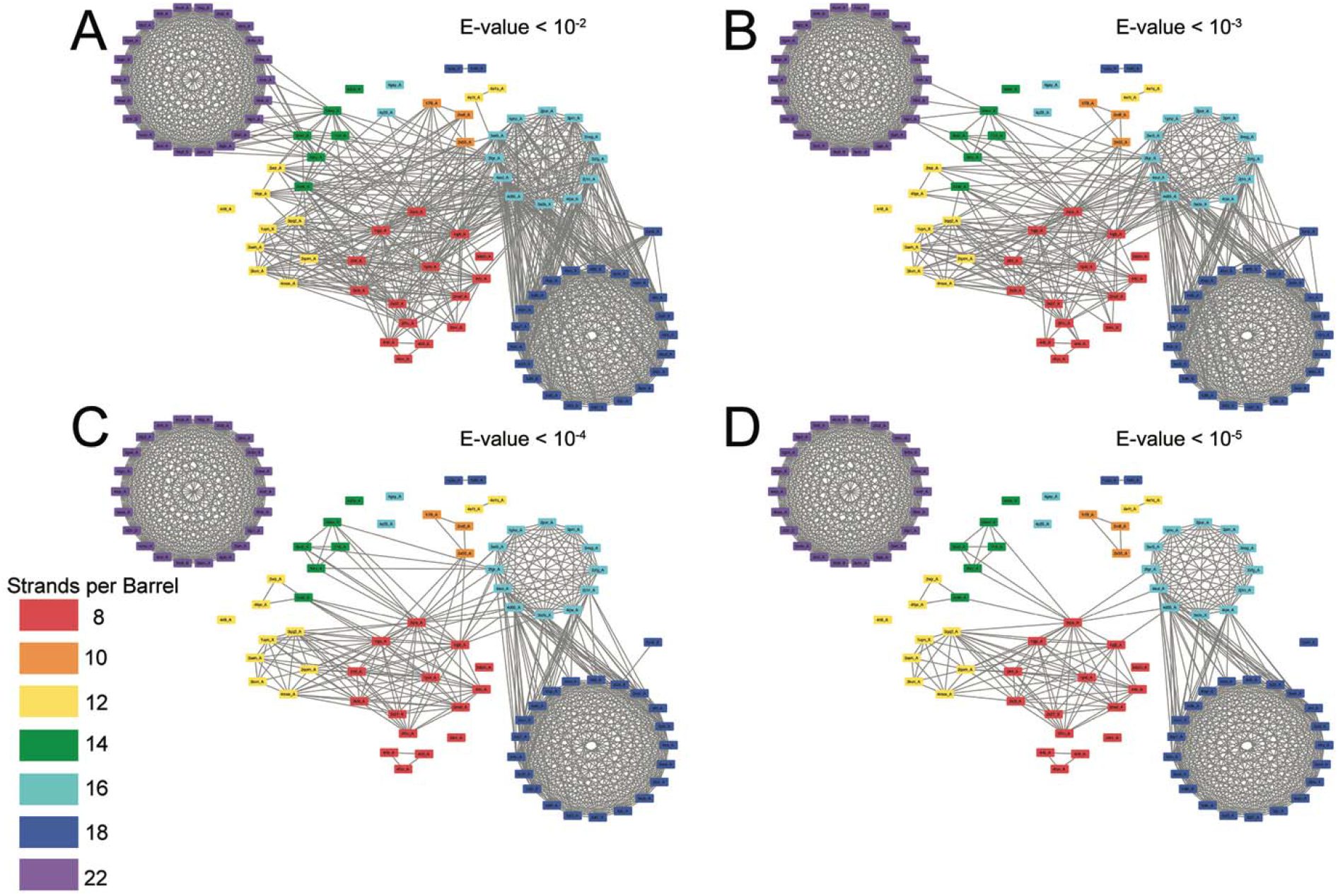
The prototypical barrels at E-values ≤ 10^−2^ to 10^−5^ colored by number of strands. Each edge represents sequence similarity between two nodes (barrels) with an E-value less than or equal to the subcaption. Multiple edges between the same pair of nodes have been removed for clarity; the edge with the lowest E-value was kept for visualization. A) E-values ≤ 10^−2^ B) E-values ≤ 10^−3^, C) E-values ≤ 10^−4^, D) E-values ≤ 10^−5^.

### Connections between and among barrels of different strand numbers

As discussed above it is generally understood that a change in strand number would be caused by an amplification event—most likely duplication—and a change in sequence without a change in strand number would most likely be caused by the accretion of mutations. In order to assess the relative frequency of diversifying and then fixing amplification events versus mutation events, we compared the frequency of finding an alignment between barrels of different strand numbers to alignments of barrels with the same strand number. We assessed both the quantity and quality of the alignments between different and among same strand numbers. In order to not over preference the frequency and quality of the alignments among barrels of the same size we narrowed our dataset to the 55 barrels of only 25% sequence similarity; an equivalent of fig. 1D at 25% sequence similarity is shown in fig S1.

With respect to the quality of alignments, E-values are almost always lower among barrels of the same strand number than between barrels of different strand number (Table S2). The only exception to this is the 8-stranded barrels; this group has slightly lower average E-values with the two 10-stranded barrels than they do among themselves. 10-, 12-, 14, 16-, 18-, and 22-stranded barrels all have lower E-values for the alignments among barrels of the same strand number than between barrels of different strand numbers.

**Figure 2.**
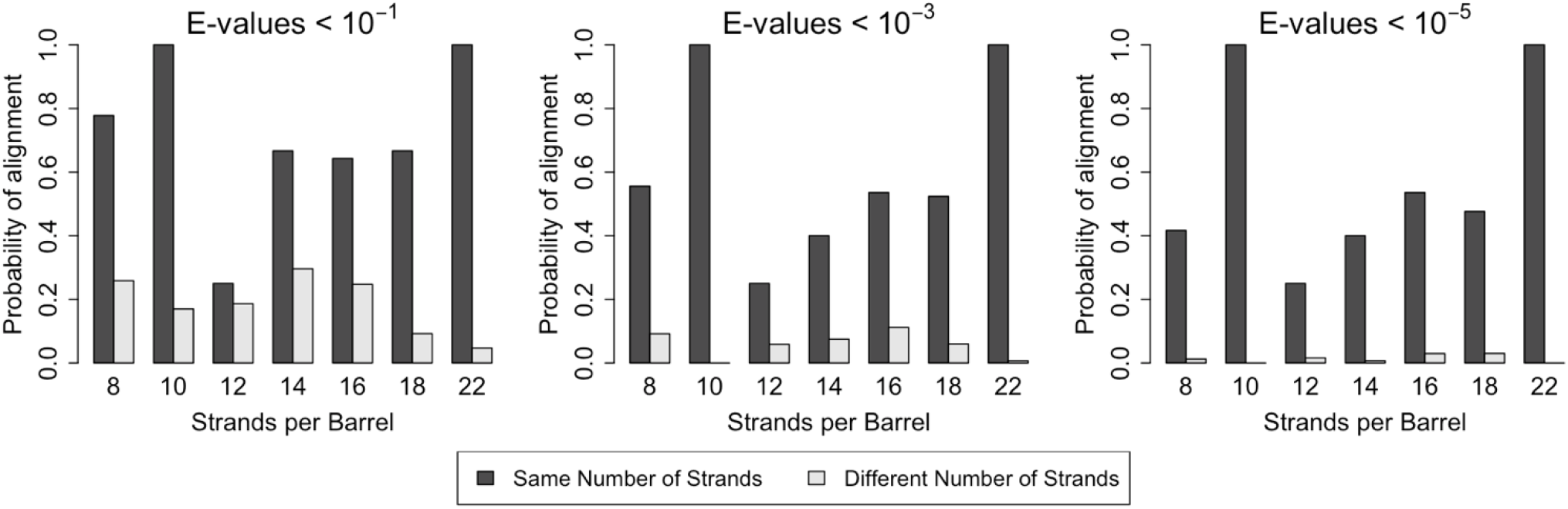
Similarity is particularly high among prototypical barrels with the same number of strands. Frequency that these barrels aligning with other prototypical barrels of the same number of strands compared to the frequency these barrels aligning with prototypical barrels with different numbers of strands. Only prototypical OMBBs with ≥25% sequence similarity were considered. Alignments of OMBBs shown at E-values < 10^−1^ (left), E-values < 10^3^ (center), E-values < 10^−5^ (right). If multiple alignments between two proteins existed, we used only the alignment with the smallest E-value. Probability was calculated as the number of alignments in each group divided by the possible combinations of barrel alignments for each group.

With respect to quantity of alignments, there is, in all cases, a higher probability of alignments between strands of the same size barrel than alignments between different sized barrels (fig. 2). Although duplication events are known to be more frequent than mutation events, the high frequency and high quality of the alignments between barrels of the same size strongly suggest that for OMBBs mutation events are much more frequently fixed than duplication events.

### Internal Repeats

For all barrel types delineated above, in order to better understand the repeat nature of the evolution from one strand number to another, alignments of sequence similar strands can be found internally by allowing the same barrel to self-align, just as they were done externally by aligning different barrels with each other. We will refer to the internal alignments as internal repeats and external alignments as alignments or conserved residues.

All internal repeats within prototypical barrels are hairpin shifts or double hairpin shifts. A hairpin shift is where a set of strands aligns in two ways and the first two strands of the two repeats are shifted over by two strands i.e. a hairpin; for example, a 7-stranded hairpin shift would be an alignment between strands 1-7 and strands 3-9. A double hairpin shift is where a set of strands aligns in two ways and the aligned strands of the two alignments are shifted over by four strands from each other.

Internal repeats are more characteristic of some beta barrels than others (fig. 3A). Specifically, they are more common for the smaller barrels than the larger ones. Although a full register of the internal alignments for all 130 proteins can be found in the supplement (InternalRepeatsE10-3.xlsx), here we enumerate the patterns that we find in the prototypical barrels (fig. 3B).

The 12-, 14-, 18-, and 22-stranded barrels have single hairpin shifts. The 8- and 16-, stranded barrels sometimes have double hairpin shifts (8/20 repeats) and sometimes have single hairpin shifts (12/20 repeats) (fig. 3C and 3D). The only 10-stranded barrel with an internal repeat has a double hairpin shift. It is unclear how these repeats came about. Regardless of the internal repeats that do exist, the 12-, 14-, 16-, and 18-stranded barrels all contain at least one strand to which no internal repeat could be identified. In the 16- and 18-stranded barrels there is rarely an internal repeat alignment that includes the first strand (19/20 cases do not have it).

**Figure 3.**
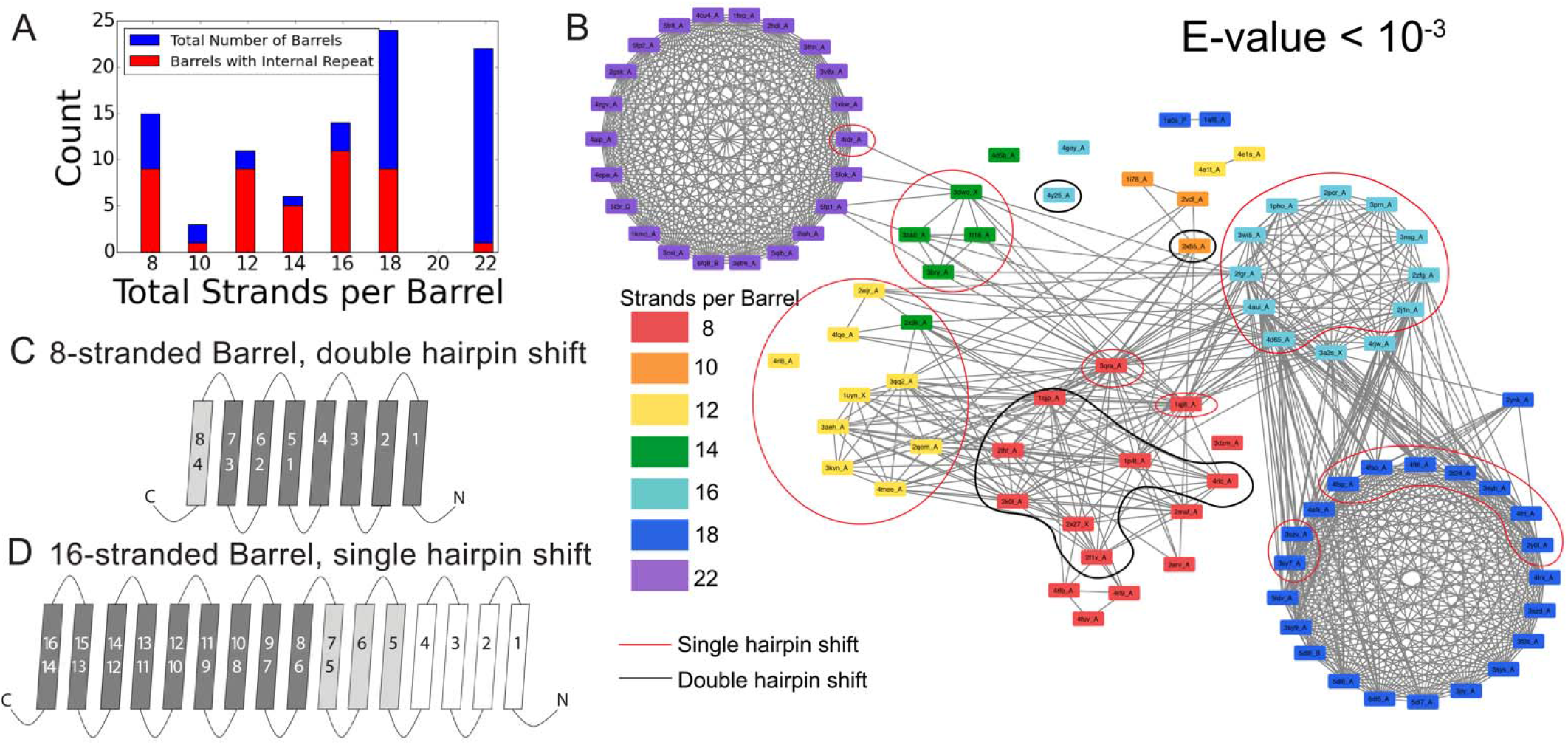
Internal repeats of the well-connected prototypical barrels. A) Distribution of all prototypical barrels with an internal repeat. The blue bars represent the total counts, while the red bars represent the proportion with an internal repeat. B) Internal repeats are identified for the barrels that are circled. Red circles indicate a single hairpin shift; black indicates a double hairpin shift (See text for definition). C,D) Strands are shown going right to left, N to C, following the orientation in the membrane, with up indicating extracellular and down indicating intracellular. Dark grey strands indicate that all barrels of that size share the same repeat pattern; light grey strands indicate that only a few of the barrels have that extension to the internal repeat. White strands indicate that these strands are never observed to participate in an internal repeat. The top row of numbers represents the strand number and the bottom row of numbers represents the strand numbers that align with the top numbers in the internal repeat C) 8-stranded barrels, D) 16-stranded barrels.

### Alignments

Internal repeats show the duplications that lead to a full-length protein while alignments between proteins can tell us how the proteins are evolutionarily linked to each other.

A potential evolutionary pathway emerges when considering the alignments between barrels of different strand numbers in the prototypical group (fig. 4). Here we describe these alignments in two steps, how the eight-stranded barrels align with the 10-, 12-, 14-, and 16-stranded barrels and then how 14-stranded barrels align with 22-stranded barrels and how 16-stranded barrels align with 18-stranded barrels. The same strands that align within the first step also align in the second step. So although 8-stranded barrels do not align with 18- or 22- stranded barrels, the strands are passed through. This means that the strands from 8-stranded barrels that align with the 16-stranded barrels are the strands in the 16-stranded barrels that align with the 18-stranded barrels. Similarly, the strands in the 8-stranded barrels that align with 14-stranded barrel are the strands in the 14-stranded barrel that align with the 22-stranded barrels.

Alignments discussed below in step 1 and step 2 are all at an E-value≤10^−2^. Examples of sequence alignments between barrels of different sizes are shown in the supplement (fig. S2).

#### Step 1

The diversification from barrels with 8 strands to 10-, 12-, 14-, and 16-stranded barrels is the first step of strand number diversification. The eight-stranded barrels are at the center of the network and have similar alignments to 10-, 12-, and 16-stranded barrels (fig. 1A-C). In particular, when the eight-stranded barrels align with the other barrels, the alignment includes at least two and often as many as all eight (fig. S2, fig. S3A) of the strands. This manifests as the 8-stranded barrels often (139/152 cases) aligning with the last eight strands of the 10-, 12-, and 16-stranded barrels such that the last strands align, the penultimate strands align, the third-to-last strands align, etc. (top step of fig. 4). However, all 8 strands of the 8-stranded barrel are not always involved in these alignments-just 44/139 cases of this type of alignment contain 8 strands. Yet the position of the alignment does not change. All 139 alignments contain the penultimate strands.

The alignment between 8-stranded barrels and 14-stranded barrels is more complex. There are two major patterns of alignments between the eight-stranded barrels and the 14-stranded barrels. The first is a seven-stranded unit encompassing the last seven strands of both barrels (22/36 cases). The second is a seven-stranded unit with a hairpin shift from the first alignment (10/36 cases). This alignment is therefore between strands 2-8 and strands 6-12 of the 8- and 14-stranded barrels respectively (fig. 4). These alignments are between 3 and 8 strands long (fig. S3D).

**Figure 4.**
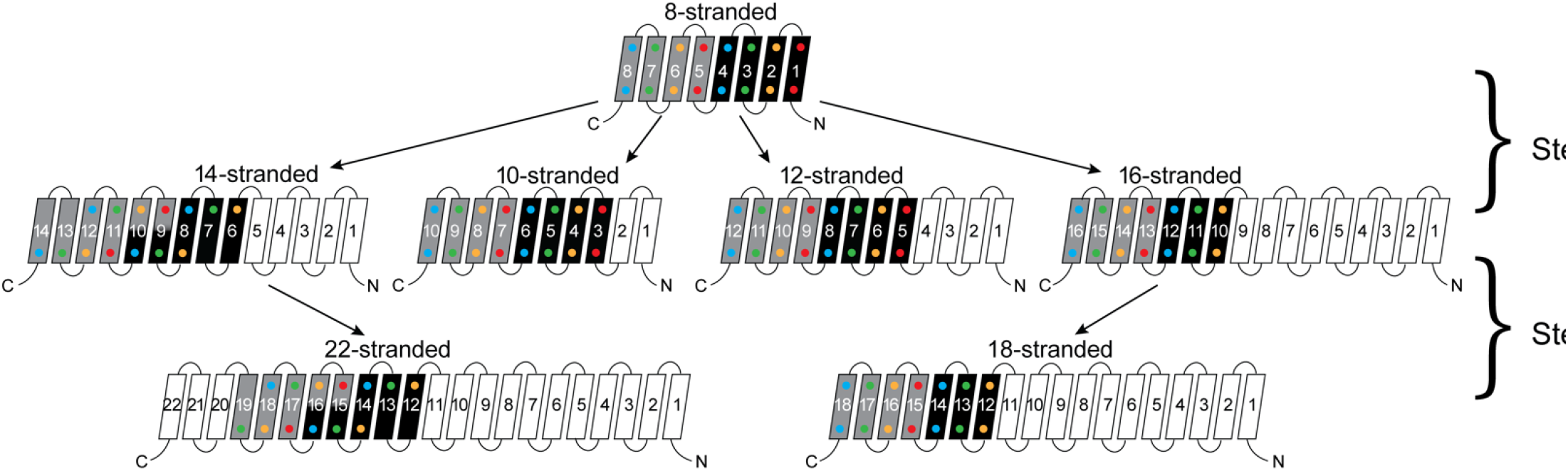
Persistence of the 8 strands in strand diversification from 8 to 22 strands. 8-stranded proteins are found to align to proteins with 10, 12, 14 and 16 strands. The strands that initially align from the 8-stranded barrels in the 16-stranded barrels can be aligned from 16-stranded barrels to 18-stranded barre, Moreover, the strands that align between the 8- and 14-stranded barrels align between the 14- and 22- stranded barrels. However 8-straded barrels themselves do not directly align with the 18- or 22- stranded barrels. Arrows indicate alignment between proteins of different strand numbers. In 8-stranded barrels (but only in 8-stranded barrels), black strands have an internal repeat with gray strands. Colored dots (red, yellow, green and blue) indicate which strands are aligned, i.e. black with red dots to black with red dots, gray with yellow dots to gray with yellow dots. In the case of hairpin shifts for 14- and 22-stranded barrels, the two alternative alignments are depicted by split strands with different backgrounds and dot colors. For example, in 14-stranded barrels strand nine aligns with strand three and strand 5 from the 8-stranded barre and is consequently colored half like strand three and half like strand 5. White strands are strands that are not found to align to the original eight strands between barrels of different sizes.

#### Step 2

The diversification from 14 to 22 strands and from 16 to 18 strands is the second step of the strand number diversification. The alignment from the 14- to the 22-stranded barrel is relatively simple (fig. S3E). All alignments are between strands 4-13 of the 14-stranded barrels and strands 10-19 of the 22-stranded barrels.

The alignments from the 16-stranded barrels to 18-stranded barrel are the most complex (fig. S3F). In most alignments, the 16-stranded barrel aligns with either the first 16 or last 16 strands of the 18-stranded barrel, with two additional strands added to form the full 18-stranded barrel; shorter alignments that are subsets of either of these two alignments are also commonly observed. 215 of 225 alignments between the 16- and 18-stranded barrels fall into these two categories. However, there are a few notable exceptions (fig. 5).

**Figure 5.**
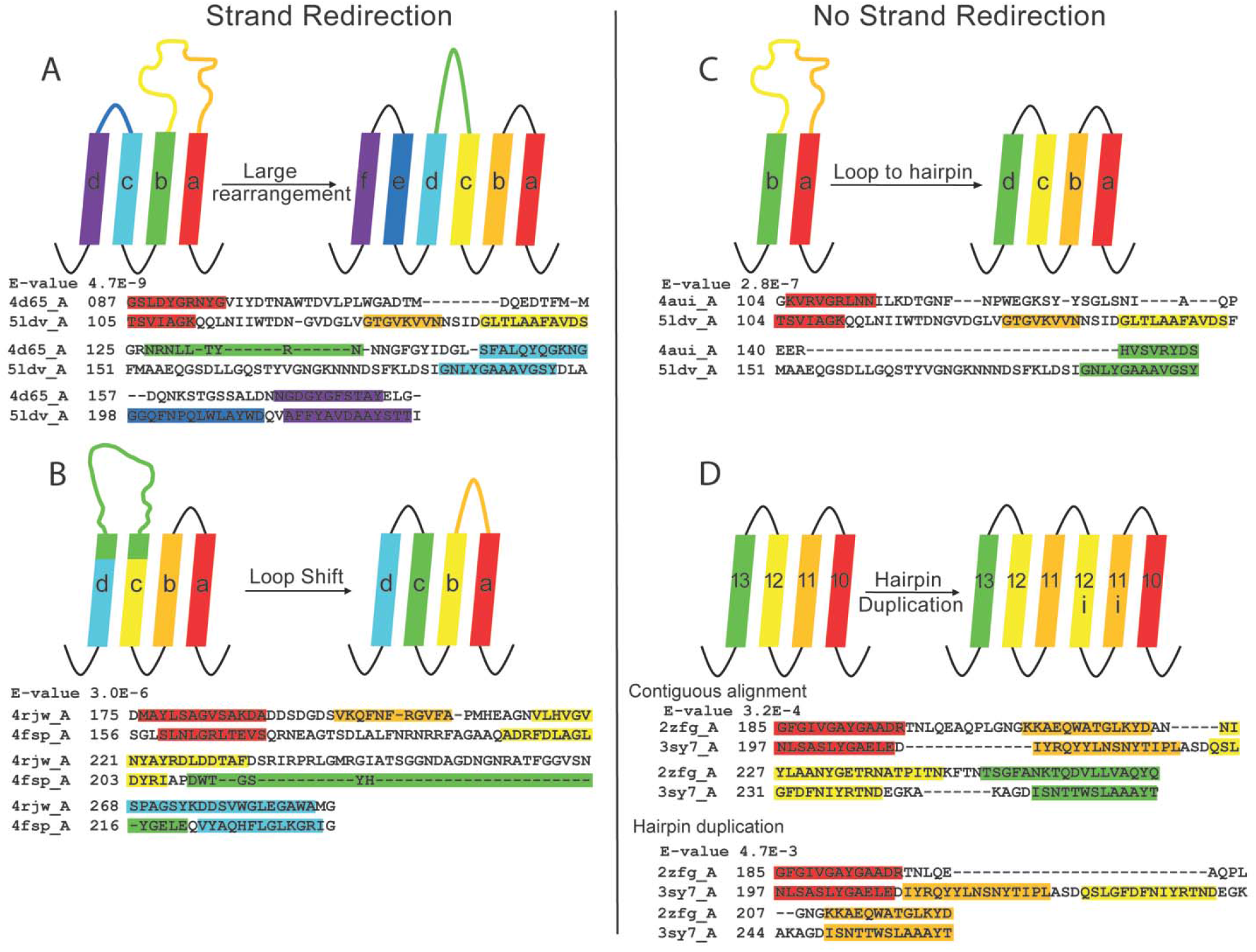
The unusual alignments of the 16- to 18-stranded barrels. In all panels, the 16-stranded barrel is shown on the left; the 18-stranded barrel to which it is aligned is on the right. Loops are colored to reflect the strands that they were or have become. Strand boundaries are determined by a combination of phi psi angles and hydrogen bonding patterns as described in the methods. A representative alignment is shown below each diagram. Only strands are colored in the alignments to match the colors in the diagrams. A) Large rearrangement. In this rearrangement, the loop between a and b in the 16-stranded barrel becomes two strands in the 18-stranded barrel and strand b of the 16-stranded barrel becomes a loop in the 18-stranded barrel. B) Loop shift. The long orange colored loop of the 16-stranded barrel shifts to form a strand in the 18-stranded barrel, this redirects the yellow strand in the 18-stranded barrel, while parts of two other strands (green) of the 16-stranded barrel create a new strand thereby preventing further strand misdirection. C) Loop to hairpin. A large loop in the 16-stranded barrel (yellow/orange) becomes two strands in the 18-stranded barrel. D) Hairpin duplication. A hairpin from the 16-stranded barrel (yellow/orange) is duplicated in the 18-stranded barrel.

These exceptions are perhaps the most interesting as they illustrate possible new, non-duplication methods of strand number diversification. These alignment exceptions show proteins that do not map linearly from the strands of 16-stranded barrel to the strands of the 18-stranded barrel. There are four mechanisms for the different forms of alignment that occur between 16-stranded barrels and 18-stranded barrels. These mechanisms were discovered by analyzing sequence alignments with the crystal structures. Full alignments with E-values are shown in supplement fig S4.

The first mechanism is a large rearrangement in which three loops and two strands in the 16-stranded barrels are rearranged to form five loops and four strands (fig. 5A). The very long loop of the 16-stranded barrels folds into a hairpin, displacing the next strand to a loop; the third loop also moves into a strand conformation. This has the unusual effect of aligning an extracellular-facing strand to a periplasmic-facing strand. The cyan strand on the left is extracellular-facing, but it aligns to the periplasmic-facing strand on the right. We observe two instances of this type of rearrangement. In these two cases, two 16-stranded proteins (PDB ID 2FGR and 4D65) align to create the same large rearrangement with the same 18-stranded barrel (PDB ID 5LDV). It is unlikely that both proteins made the same unusual large rearrangement. More likely is that since the two 16-stranded barrels are highly similar, when one evolved a large rearrangement to an 18-stranded barrel the other 16-stranded barrel maintained a strong alignment with the newly formed 18-stranded barrel.

Beta-strands have a sequence hallmark of alternating polar and non-polar amino acids. Therefore, we checked for an increase in polarity alternation between the sequences that were loops in the 16-stranded barrels which became strands in the 18-stranded barrels and the strand in the 16-stranded barrel that became a loop in the 18-stranded barrel. The strands that were loops average 58% alternation and the loops that were strands average 51% alternation. However, it is difficult to determine if this increase of alternation in strands over the loops they are aligned with is statistically significant for such few counts.

The second type of rearrangement is a loop shift (fig. 5B); one loop becomes a strand and a nearby strand becomes a loop. This also has the effect of aligning periplasmic- and extracellular-facing strands. Overall, this does not change the number of strands in the barrel, but it does influence strand directionality and the location of the long loops. We observe four instances of this type of rearrangement, from a single 16-stranded barrel (PDB ID 4RJW) to four different 18-stranded barrels (PDB ID 2Y0L, 3SY9, 4FSP, 4FT6). A single 16-stranded barrel aligning in this way with four 18-stranded barrels would strongly implicate a single 16- to 18- stranded diversification event and then subsequent diversification among the 18-stranded barrels.

Finally, there are two rearrangements that do not result in strand redirection. We observe just one example of a loop to hairpin transition (fig. 5C) in which a long loop folds down to form a new hairpin between two existing strands. This occurs for an 18-stranded barrel where we also see an alignment of a large rearrangement as described above (PDBID 4AUI to 5LDV). It is likely that only one of these diversification events actually occurred and the other is a residual alignment to other closely related barrels. In the case of the large rearrangement there is a lower E-value than in the loop to hairpin alignment though that does not mean that the large rearrangement is necessarily the true mechanism. Similar to the analysis of the large rearrangement, the loops have an average of a 46% strand alternation and the strands they become have an average of a 59% alternation. Determining statistical significance for these small counts is not possible.

The hairpin duplication (fig. 5D) adds an extra copy of the hairpin formed by strands 11 and 12; in the alignments, this appears to be similar to the loop to hairpin conversion. However, in all three examples of hairpin duplication, there exists a long gap in the 16-stranded barrel where no residues can be aligned. These three alignments occur for one 16-stranded barrel with three different 18-stranded barrels (PDB ID 2ZFG to 3SY7, 3SZV, and 4FT6) (fig. 5D, bottom alignment). It should be noted that the 18-stranded barrel in PDB ID 4FT6 is also seen as a possible loop shift mechanism with a lower E-value.

Additionally, for each of the hairpin duplication alignments there is an alternative alignment in which the duplication does not occur and strands 11 and 12 align in both barrels (fig. 5D, top alignment). This suggests that instead of the hairpin duplicating, there is a C-terminal hairpin that is added, similar to the most common form of 16-stranded to 18-stranded alignment.

Both the loop to hairpin and the hairpin duplication rearrangements influence the total number of strands, but no misdirection of strand topology occurs. Although we searched for misdirections of strand directions in any other alignments of barrels of other sizes, we found no other examples outside the 16- to 18-stranded alignments described.

### C-terminal conservation

As described above barrels of the same strand number are better aligned than barrels of different strand number. The patterns of barrel alignments allow us to trace a conserved, possibly “original”, set of strands through all the barrel sizes (fig. 4). The last seven strands of the 8-stranded barrels can be traced to the last strands of the 10-, 12-, 14-, 16- and 18-stranded barrels; they are also similar to the middle of the 22-stranded barrels. In the case of the 14- stranded barrels, there are two alternate alignments to the 8-stranded barrels; this is reflected by the two-tone pattern to the 14-stranded barrel. As indicated by the tree-like pattern in fig. 4, most of these connections are not direct and must be followed through two separate alignments. For example, the 8-stranded barrels align to the 16-stranded barrels, and the 16-stranded barrels align to the 18-stranded barrels, but the 8-stranded barrels do not align to the 18-stranded barrels.

**Figure 6.**
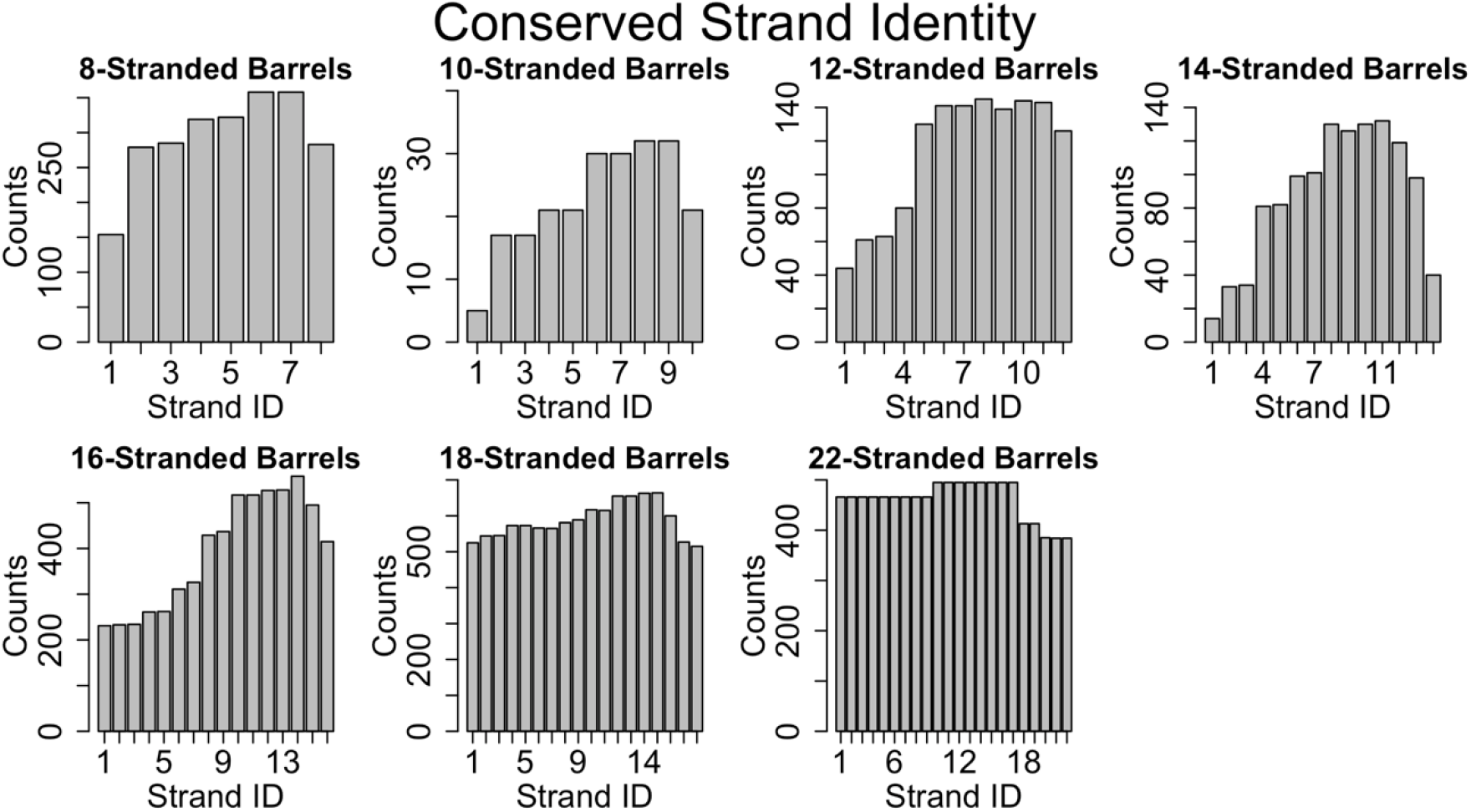
Conservation of strands in the prototypical barrels at <85% sequence identity. For each alignment involving two prototypical barrels, the identity of the strands in each barrel was determined. Each graph represents the distribution of the strands reused in that barrel size.

Documenting these repeats led to the observation that we find a significant conservation of strands in the C-terminal half of the protein and a much lower conservation overall for the N-terminus (fig. 6, fig. S5 at 25% sequence identity shows a similar trend). Specifically, by conservation we mean reuse of sequence between proteins. We find this C-terminal half conservation to be the case for proteins of all strand numbers.

## Discussion

Looking at a combination of structural and sequence information allows us to better understand the evolutionary relationships among outer membrane proteins. Grouping outer membrane proteins by sequence relationship results in groups that are united by function and have structural similarities. Combining sequence and structural information allows us to trace evolutionary pathways among these proteins.

### Prevalence of divergence from mutations vs duplication

Our analysis of the relative frequency and quality of alignments between barrels of the same size and barrels of different sizes illustrate that alignments are always more likely to be within the same strand number than between barrels of different strand numbers (Fig 2, Table S2). Moreover as demonstrated by lower E-values, the quality of the alignments among barrels of the same strand numbers are better than the quality of alignments between barrels of different strand numbers. This demonstrates that although it is known overall that duplication events are more common than mutation events, for OMBBs partial-gene duplication events that are fixed are rare. We conclude from this that most diversification among OMBBs occurs through mutation and that events that change the strand number are rare. This bolsters understanding of strand number changes as discrete events which can be charted as steps of diversification.

### Internal repeats and the primordial barrel

The large percentage of prototypical barrels that have internal alignments is highly suggestive that strand number diversification - when it occurs—largely results from amplification. Our data is consistent with these duplications occurring for many different numbers of strands. Although the 8-stranded barrel is the smallest known OMBB single chain barrel, we see repeats of four strands within the 8-stranded barrels. This points to the idea that the 8-stranded beta barrel evolved from a smaller segment, likely the double hairpin described previously (Remmert, et al. 2010). Given the large number of 3- and 7-stranded alignments (fig. S6), this also suggests that the first or last strand of these double hairpin units has degenerated, possibly due to evolutionary pressure associated with needing to interact with other strands.

A primordial 8-stranded barrel with an internal double hairpin repeat gives rise to the possibility that a primordial four strands formed a multi-chain beta barrel. These stands would then ultimately be duplicated for the first single-chain barrel. Since most repeat proteins do not have function as individual domains, it has been hypothesized that many repeat proteins were originally homo-oligomers (Ponting and Russell 2000). This hypothesis suggests that for beta barrels the primordial barrel was a multi-chain beta barrel like YadA, a trimeric adhesion protein, where each chain contributes four strands (a double hairpin) to create a 12-stranded barrel (Franklin, et al. 2018).

### A pathway for strand number diversification

Our graph of strand number connectivity (fig. 3D) demonstrates a likely evolutionary path towards strand number diversification. The alignment data is consistent with the notion of a primordial 8-stranded barrel that developed more strands at the N-terminus through hairpin duplication, or some evolutionary mechanism to ultimately create 10-, 12-, 14-, and 16-stranded barrels. From these branching points, there were two secondary evolutionary transitions with 22-stranded barrels evolving from the 14-stranded barrels and 18-stranded barrels evolving from the 16-stranded barrels.

The consistent location of the strands that are aligned suggests that either there was a single event that caused the additional strands and then diversification from that point, or that there is a preferred pathway of duplication in which all strand increase events must participate. The fact that the 18- and 22-stranded barrels overwhelmingly have alignments among them that encompass the entirety of both barrels (fig. 6 and fig. S6) suggests the possibility that barrels of these sizes are all orthologs.

### Novel diversification mechanisms

The pattern of conservation in the 16- to 18-stranded barrels illustrates that a repeating topology can be created without motif duplications. The alignments between 16- and 18-stranded barrels suggest more than one path to an addition of two strands. Since we are hypothesizing that the 16-strand to 18-strand barrel transition event is among the more recent of the strand number diversification events (i.e. step 2), it makes sense that we can map these events more clearly than the other strand diversification events. These novel diversification mechanisms may have been used by other strand number transitions, but since those transitions may be older they would be more likely to have been obscured by further evolution.

Some of the transitions between 16- and 18-stranded barrels are through large rearrangements and shifts that used existing amino acids to generate the additional two new strands rather than generating them through gene duplication (fig. 5). The types of rearrangements we see imply that the boundary between strand and loop may be somewhat fluid. Since half of strand residues are hydrophilic, membrane bound strands aren’t too hydrophobic to become loops. Conversely loops can become strands. The accretion of a few localized polar amino acids may cause a tight turn to be more favorable and insertion into the membrane less favorable.

The novel mechanisms of diversification of strand number are a strong counter argument to the intuitive understanding that all repeat segments in repeat proteins are made from gene/motif duplications. Moreover, that a repeat segment is created by something other than a duplication event is suggestive that the environment and function required the repeated structural motif. In the process of collecting this data, other iterations of the alignment dataset were created with slightly fewer sequence inputs. These generated different alignments, but the same novel mechanisms of diversification were present. Therefore, it appears that the environment and function of the barrel enforce the structural repeats of hairpins in the beta barrel even without sequence repeats.

### No change in direction

The existence of strand number diversification events that are not hairpin duplications means that it could also be single strands added through diversification events. If single strand additions are possible, why are all bacterial OMBBs even stranded? We believe that this is because the strands and loops encode a membrane topology that is difficult to override. Unlike inner membrane proteins with a positive inside rule (von Heijne and Gavel 1988), outer membranes show a charge out rule within their strands (Seshadri, et al. 1998; Slusky and Dunbrack 2013). Addition of an odd number of strands anywhere but the C-terminus would change the directionality of all subsequent strands. Preference for directional maintenance is so intense that of all 2974 alignments represented in Figure 1, there are just six instances of strand redirection (fig. 5A, 5B) in which any strands change topology (from N-terminus periplasmic to N-terminus extracellular or vice versa). The six instances we document (fig. 5A, 5B) all have a strand that flips from the N-terminal end of the strand facing into the cell to the N-terminal end of the strand facing out of the cell. The topological rigidity of beta barrels contrasts with helical proteins which have documented examples of whole proteins which exist and function in both topologies (Rapp, et al. 2006).

### Increasing strand numbers more prevalent than decreasing

The relational network shows frequent strand addition and few strand subtractions. Viewing figure 1 in light of the described concept of a primordial 8 stranded barrel (figure 4), the only clear loss of strands is the two 12-stranded barrels branching off of the 14-stranded 2×9k. The lack of more obvious strand loss can be attributed to either one or a combination of three causes: 1) The rate of gene or domain deletions in bacteria (Nilsson, et al. 2005) is generally regarded to be lower than that of gene or domain duplications in bacteria (Starlinger 1977; Reams, et al. 2010; Katju and Bergthorsson 2013); 2) Because the primordial single-chain barrel appears to be the smallest, i.e. 8-stranded, starting from the primordial barrel would require two strand change events—one addition and then one subtraction—in order to see a subtraction; 3) Strand change events are sufficiently difficult to fix and happen infrequently enough that the sample size for such measurements is too small.

### No 20-stranded barrels

It is somewhat surprising that structures have been resolved for every even number of stranded barrel from 8 to 26 excluding 20-stranded barrels (fig 3A). Even the more permissive search used by Reddy and Saier which predicts over 50,000 barrels does not find any predicted 20-stranded OMBBs for bacteria (Reddy and Saier 2016).

Our sequence similarity networks shed some light on this mystery. Clearly, OMBB strand numbers are not randomly distributed. Moreover, though through evolution amplification events might be common, we see very few of them fixed. Strand number transitions are rare, likely requiring distinct and improbable events. Based on the type of transitions we have seen, the most likely pathways would be a doubling of a 10-stranded barrel or one of the mechanisms of strand addition seen in the 16 to 18 strand transition applied to an 18-stranded barrel. However, 10-stranded barrels are rare enough themselves with only three occurrences in our data set. Moreover, a strand addition from 18 would be three transitions away from the primordial group which is a number of steps that has not yet been documented. Given the rarity of amplification events it may be that 20-stranded barrels have not yet been sampled.

### C-terminal strand conservation

Our results demonstrate that although it is known that membrane proteins diversify more than other proteins (Shimizu, et al. 2004; Sojo, et al. 2016), the diversification is applied unequally. It is easier to find evolutionary traces in the C-terminal half of outer membrane proteins than in the N-terminal half. This is most distinct for 16 stranded barrels but can be seen as a feature of OMBBs more broadly (fig. 6, fig S5). These results are either due to negative selection, i.e., suppression of fixation in the C-terminal half, or positive selection that leads to diversification in the N-terminal half.

The importance of the strands on the C-terminal half of the protein has been observed before. For example, mutational studies of 8-stranded OmpA identified strand 6 as being particularly important for folding (Koebnik 1999; Stapleton, et al. 2015). Duplication studies of OmpX also successfully folded, as long as the strands of the middle hairpin (strands 4 and 5) were included (Arnold, et al. 2007). This may explain unsuccessful attempts as well; to create a 16-stranded barrel by adding to the C-terminus, the last hairpin was duplicated in the 14-stranded OmpG but IR studies revealed that this additional hairpin never formed as part of the barrel (Korkmaz, et al. 2015).

The conservation in the C-terminal half does not appear to be related to the beta signal sequence (Struyve, et al. 1991) which would be located in the final strand of the barrel as the final strand appears less conserved. Rather, this pattern of reuse may be a result of (post-translational) folding the C-terminal half of the barrel first, as has recently been posited (Maurya and Mahalakshmi 2016). If this is shown to be true for other proteins, it may suggest that C-terminal conservation protects the folding pathway. Previous investigation of soluble proteins has shown that the folding nucleus of proteins is more conserved in order to preserve the ability of the protein to fold (Mirny and Shakhnovich 2001). The C-terminal half may nucleate folding by inserting the first beta-strands thereby creating a template for further beta-strand formation (Figure 7A). Alternately, this C-terminal half conservation also supports the idea that the folding mechanism might be similar to the mechanism recently proposed for the mitochondrial SAM50 (Höhr, et al. 2018). In this scenario BamA would incorporate C-terminal strands of the unfolded client protein into the assembly barrel with the N terminus of the client protein being held in the pore of the assembly protein (Fig 7B). The folding mechanism of more bacterial prototypical beta barrels will need to be assessed before the reason for conservation of the strands on the C-terminal half of the protein can be fully resolved.

**Figure 7.**
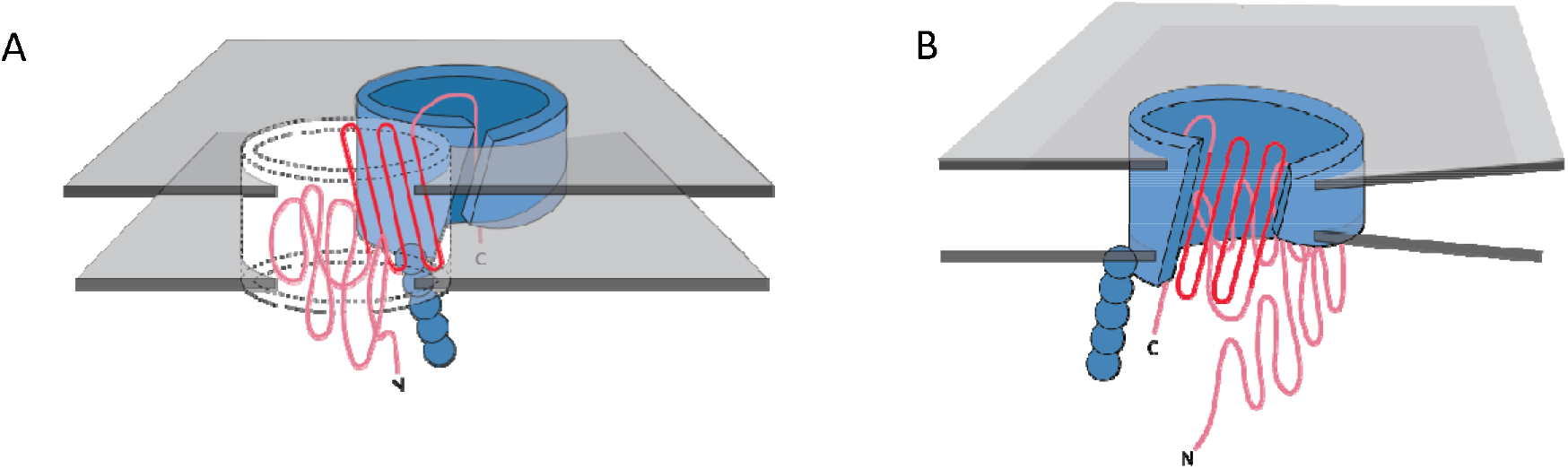
Possible mechanisms of OMBB folding. The C-terminal half of beta barrels evince a strong evolutionary trace. **A. Nucleation**: The strong evolutionary trace may be indicative of nucleation. Such nucleation could occur with the second half of the nascent barrel (red squiggly lines) forming beta hairpins in the membrane (boundaries shown with gray planes) along strands of the outer membrane translocation machinery, BamA with POTRA domains (Solid blue barrel with five attached spheres). The rest of the nascent barrel would use the formed strands as a template. **B. Incorporation**: the strong evolutionary trace may also be indicative of beta strand incorporation, whereby the second half of the beta barrel is incorporated into BamA. In this model the conserved strands form new strands in the BamA barrel while the N-terminal half of the protein inserts through the center of the barrel. Ultimately the nascent barrel buds off of the BamA, creating a new OMBB.

Overall, mapping sequence alignments with solved structures of prototypical beta barrels sheds light on the evolution of these diverse proteins. As greater numbers of prototypical beta barrels are structurally characterized more links will be elucidated. However, even with many more structurally resolved beta barrels, we may never see the branches that fully map the diversification space as steps are swallowed by evolution.

## Methods

### Barrel Network Creation

The barrel network was created as described previously (Franklin, et al. 2018). Briefly, we identified evolutionary relationships using the sensitive HMM sequence aligner HHSearch (Soding 2005), for the set of 130 beta barrels which are 85% sequence similar. HMM profiles for the 57 beta barrels in our list but not in this database were generated using the webserver for HHblits (Remmert, et al. 2012) and the database uniclust30_2017_04. The prototypical barrel cluster was defined as the large cluster that had no connection with an E-value < 1 to any other OMBB.

We organized the set of proteins and their alignments as a network, using CyToScape (Saito, et al. 2012), and CyToStruct (Nepomnyachiy, et al. 2015) to easily view the alignments in a molecular viewer. The resulting network is available online at: http://cytostruct.info/rachel/barrels/. Users can change the E-value cutoff and see the resulting networks: using a more stringent cutoff removes edges, resulting in a more fragmented network. The structure of a protein can be viewed in a molecular viewer by clicking its node. The structural and sequence alignment can be viewed by clicking the edge.

### Internal Repeats

Internal repeats were identified by self-aligning each of the 130 beta barrels using HHalign. Results were filtered to exclude alignments with an HHalign probability >75% and E-value≤10^−3^. The HHalign probability is the log-odds probability of whether the alignment was produced by the HMM versus a random model (Soding 2005). The full set of results can be found in the supplemental file InternalRepeatsE10-3.xlsx.

## Acknowledgements

Professors James Walters, Mark Holder, Christian Ray and Eric Deeds for helpful discussions and Amy Weiss for illustrations. NIH awards DP2GM128201, P20GM103418, P20GM113117, T32-GM008359, NSF MCB160205, The Gordon and Betty Moore Inventor Fellowship, KU-startup and Israel Science Foundation grant number 450/16.

## References

Arnold T, Poynor M, Nussberger S, Lupas AN, Linke D. 2007. Gene Duplication of the Eight-stranded β-Barrel OmpX Produces a Functional Pore: A Scenario for the Evolution of Transmembrane β-Barrels. Journal of Molecular Biology 366:1174-1184.

Dong C, Beis K, Nesper J, Brunkan-LaMontagne AL, Clarke BR, Whitfield C, Naismith JH. 2006. Wza the translocon for E. coli capsular polysaccharides defines a new class of membrane protein. Nature 444:226-229.

Drake JW, Charlesworth B, Charlesworth D, Crow JF. 1998. Rates of Spontaneous Mutation. Genetics 148:1667.

Franklin MW, Nepomnyachiy S, Feehan R, Ben-Tal N, Kolodny R, Slusky JSG. 2018. Efflux Pumps Represent Possible Evolutionary Convergence onto the Beta Barrel Fold. BioRxiv.

Höhr AIC, Lindau C, Wirth C, Qiu J, Stroud DA, Kutik S, Guiard B, Hunte C, Becker T, Pfanner N, et al. 2018. Membrane protein insertion through a mitochondrial β-barrel gate. Science 359.

Jimenez-Morales D, Liang J. 2011. Pattern of Amino Acid Substitutions in Transmembrane Domains of β-Barrel Membrane Proteins for Detecting Remote Homologs in Bacteria and Mitochondria. PLoS ONE 6:e26400.

Katju V, Bergthorsson U. 2013. Copy-number changes in evolution: rates, fitness effects and adaptive significance. Frontiers in Genetics 4.

Koebnik R. 1999. Membrane assembly of the Escherichia coli outer membrane protein OmpA: exploring sequence constraints on transmembrane [beta]-strands. Journal of Molecular Biology 285:1801-1810.

Korkmaz F, van Pee K, Yildiz Ö. 2015. IR-spectroscopic characterization of an elongated OmpG mutant. Archives of Biochemistry and Biophysics 576:73-79.

Marcotte EM, Pellegrini M, Yeates TO, Eisenberg D. 1999. A census of protein repeats 11 Edited by J. M. Thornton. Journal of Molecular Biology 293:151-160.

Maurya SR, Mahalakshmi R. 2016. Control of human VDAC-2 scaffold dynamics by interfacial tryptophans is position specific. Biochimica et Biophysica Acta 1858:2993-3004.

Mirny L, Shakhnovich E. 2001. Evolutionary conservation of the folding nucleus 11 Edited by A. R. Fersht. Journal of Molecular Biology 308:123-129.

Nepomnyachiy S, Ben-Tal N, Kolodny R. 2015. CyToStruct: Augmenting the Network Visualization of Cytoscape with the Power of Molecular Viewers. Structure 23:941-948.

Neuwald AF, Liu JS, Lawrence CE. 1995. Gibbs motif sampling: Detection of bacterial outer membrane protein repeats. Protein Science 4:1618-1632.

Nilsson Al, Koskiniemi S, Eriksson S, Kugelberg E, Hinton JCD, Andersson DI. 2005. Bacterial genome size reduction by experimental evolution. Proceedings of the National Academy of Sciences of the United States of America 102:12112-12116.

Ohno S. 1970. Evolution by Gene Duplication: Springer Berlin Heidelberg.

Pearson WR. 2013. An Introduction to Sequence Similarity (“Homology”) Searching. Current protocols in bioinformatics / editoral board, Andreas D. Baxevanis … [et al.] 03:10.1002/0471250953.bi0471250301s0471250942.

Ponting CP, Russell RB. 2000. Identification of distant homologues of fibroblast growth factors suggests a common ancestor for all β-trefoil proteins 11 Edited by J. Thornton. Journal of Molecular Biology 302:1041-1047.

Rapp M, Granseth E, Seppälä S, von HeijneG. 2006. Identification and evolution of dualtopology membrane proteins. Nature Structural & Molecular Biology 13:112.

Reams AB, Kofoid E, Savageau M, Roth JR. 2010. Duplication Frequency in a Population of Salmonella enterica Rapidly Approaches Steady State With or Without Recombination. Genetics 184:1077-1094.

Reddy BL, Saier MH, Jr. 2016. Properties and Phylogeny of 76 Families of Bacterial and Eukaryotic Organellar Outer Membrane Pore-Forming Proteins. PLoS ONE ll:e0152733.

Remmert M, Biegert A, Hauser A, Söding J. 2012. HHblits: lightning-fast iterative protein sequence searching by HMM-HMM alignment. Nature Methods 9:173-175.

Remmert M, Biegert A, Linke D, Lupas AN, Söding J. 2010. Evolution of Outer Membrane Beta-Barrels from an Ancestral Beta Beta Hairpin. Molecular Biology and Evolution 27:1348-1358.

Saito R, Smoot ME, Ono K, Ruscheinski J, Wang P-L, Lotia S, Pico AR, Bader GD, Ideker T. 2012. A travel guide to Cytoscape plugins. Nature methods 9:1069-1076.

Seshadri K, Garemyr R, Wallin E, Heijne GV, Elofsson A. 1998. Architecture of β-barrel membrane proteins: Analysis of trimeric porins. Protein Science 7:2026-2032.

Shimizu T, Mitsuke H, Noto K, Arai M. 2004. Internal Gene Duplication in the Evolution of Prokaryotic Transmembrane Proteins. Journal of Molecular Biology 339:1-15.

Slusky JSG, Dunbrack RL. 2013. Charge Asymmetry in the Proteins of the Outer Membrane. Bioinformatics 29:2122-2128.

Soding J. 2005. Protein homology detection by HMM-HMM comparison. Bioinformatics 21:951-960.

Sojo V, Dessimoz C, Pomiankowski A, Lane N. 2016. Membrane Proteins Are Dramatically Less Conserved than Water-Soluble Proteins across the Tree of Life. Molecular Biology and Evolution 33:2874-2884.

Stapleton JA, Whitehead TA, Nanda V. 2015. Computational redesign of the lipid-facing surface of the outer membrane protein OmpA. Proceedings of the National Academy of Sciences 112:9632-9637.

Starlinger P. 1977. DNA rearrangements in procaryotes. Annu Rev Genet 11:103-126.

Struyve M, Moons M, Tommassen J. 1991. Carboxy-terminal phenylalanine is essential for the correct assembly of a bacterial outer membrane protein. J Mol Biol 218:141-148.

von Heijne G, Gavel Y. 1988. Topogenic signals in integral membrane proteins. European Journal of Biochemistry 174:671-678.

Zhang J. 2003. Evolution by gene duplication: an update. Trends in Ecology & Evolution 18:292–298.

